# Successful Gene Editing of Apolipoprotein E4 to E3 in Brain of Alzheimer Model Mice After a Single IV Dose of Synthetic Exosome-Delivered CRISPR

**DOI:** 10.1101/2024.04.23.590784

**Authors:** Bruce Teter, Jesus Campagna, Chunni Zhu, Grace E. McCauley, Patricia Spilman, Donald B. Kohn, Varghese John

## Abstract

**Background:** The gene for apolipoprotein E4 (ApoE4 E4) confers an increased risk for development and lowers the age of onset of Alzheimer’s disease (AD), and is a highly suitable target for CRISPR-based editing because ApoE4 differs from ApoE3 by a single nucleotide polymorphism in the codon for residue 112 that codes for arginine (CGC) in E4 and cysteine (TGC) in E3. Editing of E4 to E3 could lower the risk of AD or ameliorate E4-related AD phenotypes. For AD, in order to deliver CRISPR components across the blood-brain barrier to the brain, we have developed a delivery platform termed ‘Synthetic Exosomes’ (SEs) – microfluidically-synthesized deformable nanovesicles approximately the size of natural exosomes that have the ability to cross the BBB and deliver cargo to the brain. Here, we describe our use of SEs carrying CRISPR to successfully edit E4 to E3 in brain tissue of an E4-expressing mouse model.

**Methods:** Several CRISPR guide RNAs (gRNA) and Cytosine Base Editor (CBE) mRNAs were synthesized by chemical and in vitro transcription syntheses, respectively. Four combinations of gRNA and CBE mRNA were tested in vitro for their relative activity to edit the E4 (cytosine) to E3 (thymine) in E4-expressing neuroblastoma (E4-N2A) and human ‘Kelly’ neuroblastoma cells, to assess which combination produced the highest E4 to E3 base editing efficiency. The CRISPR RNA combination with the highest efficiency was encapsulated in SEs and injected intravenously (IV) via the tail vein into an AD model E4-expressing (E4-5XFAD) transgenic mouse; as a negative control, an E4-5XFAD mouse was injected with empty SEs. Five days after injection, mice were euthanized and brain, liver, and buffy coat (white blood cells (WBC)) collected to determine the editing of E4 to E3 measured by Next Generation Sequencing. In addition, E3 mRNA was measured in the brain and liver and compared to the %E3 gene editing.

**Results:** The highest gRNA+CBE mRNA editing efficiency was ∼50% in E4-N2A cells and the same gRNA+CBE combination delivered in SEs to Kelly neuroblastoma cells showed 6.5% editing efficiency. In the E4-5XFAD mouse in vivo, five days after IV delivery of a single dose of the highest-activity SE-CRISPR gRNA+CBE mRNA, the percent of E4 edited to E3 was 0.14% in brain, 0.8% in liver, and 0.36% in WBCs. As evidence of functional editing, SE-CRISPR-treated mice had 0.03% E3 mRNA in brain and 0.09% E3 mRNA in liver.

**Conclusions:** While this level of ApoE4 to E3 editing achieved five days after a single IV injection of SE-CRISPR is small, it provides initial in vivo proof-of-concept that the ApoE4 gene can be successfully edited, and editing results in functional expression of ApoE3 mRNA. The findings presented herein supports further optimization of the SE-CRISPR approach to increase the level of editing in brain as part of clinical development of SE-CRISPR as a powerful novel therapeutic approach for AD.

## Introduction

The major genetic risk factor for sporadic Alzheimer’s disease (AD) is expression of apolipoprotein E4 (*APOE4*, ApoE4, E4),^1–4^ with an estimated 40-65% of people diagnosed with AD having one or two copies of the E4 gene.^5^ Homozygosity for the E4 allele confers the greatest risk for the development of AD, whereas presence of an E3 allele (heterozygous E3/E4) reduces that risk for AD.^6^ Compared to E3, E4 exacerbates the two predominant hallmarks of AD brain, Aβ amyloid plaques and tau pathology,^7–14^ and has other pleiotropic mechanisms that may be implicated in disease onset.^7–21^ Considering this association, a 2024 National Institute on Aging (NIA) working group came to a unanimous consensus that cumulative data from multiple studies in humans and animal models support that lowering E4 should be a therapeutic target for AD for E4 carriers.^22^ Current therapeutic approaches targeting E4 include immuno-neutralization to reduce E4 protein,^23^ targeting the E4 gene promoter to reduce expression levels,^24^ knocking out the E4 gene using CRISPR,^25,26^ inserting the Christchurch mutation into E4,^27^ use of chemical ‘structure correctors’ to disrupt the E4 structure,^26^ and E2 gene therapy (clinical trial NCT05400330).^28,29^

The neurotoxic effects of E4 may also be suppressed by editing the ApoE4 gene to ApoE3 or ApoE2.^30^ This approach is particularly suitable for the E4 gene, where DNA coding for E4 differs from E3 by a single nucleotide polymorphism (SNP), rs429358, within the codon for amino acid 112 which codes for arginine in E4 and cysteine in E3.^31^ Clustered regularly interspaced short palindromic repeats (CRISPR) cytosine base editing (CBE) technology, which is designed to make cytosine ’C’ to thymine ’T’ DNA substitutions ^32,33^ is ideal for editing E4 to E3 by simply changing the ’C’ in E4 to ’T’ in E3,^34–36^ and the feasibility of this method is evidenced by successful CRISPR editing E4 to E3 *in vitro* in several studies.^29,33,34,37–39^ Many of these in vitro studies show reversal of E4-specific phenotypes, for example, it has been demonstrated by Zhao *et al.* that in iPSC-derived organoids of cell lines from AD patients, ApoE4-related toxic phenotypes, including tau pathology, can largely be reversed through isogenic conversion to ApoE3 using CRISPR.^40^

CRISPR-CBE can be used to convert the ’C’ in E4 to ’T’ in E3, an irreversible DNA conversion, without requiring double-stranded DNA (dsDNA) breaks by using a Cas9 nickase (Cas9n).^41,42^ Cas9n and cytidine deaminase enzyme fusion proteins, along with endogenous DNA repair systems, are used to effect the ’C’-’T’ substitution, leading to correction of ∼15–75% of total in vitro cellular DNA with minimal (≤ 1%) INDEL formation and minimal off-target editing *vitro.*^34,41^ In addition, CBE allows editing in both dividing and non-dividing cells such as brain neurons.^43^

CRISPR base editing technology, including CBE^32,44,45^ and other CRISPR modalities ^32,44^ are being used to edit genes involved in a wide range of human diseases, including AD.^32,35,44,46–51^ The first CRISPR based therapy, CASGEVY^TM^, was recently approved by the U.S. Food and Drug Administration (FDA) for sickle cell disease,^52^ and clinical trials of gene editing for many diseases are underway,^53,54,56^, with promising phase 1 results for the liver disease transthyretin amyloidosis.^55^

Despite the promise of application of CRISPR-CBE editing for treating diseases of diverse organs, successful *in vivo* application of CRISPR technology to edit ApoE4 to ApoE3 in the brain has not been reported. This critical step in translation of E4 to E3 editing to the brain is hindered by the necessity of delivering CRISPR components across the blood-brain barrier (BBB), ultimately to nuclei of cells in the brain parenchyma, wherein editing would take place.^57^

Our lab has been developing a brain delivery platform termed ‘synthetic exosomes’ (SEs), a type of deformable lipid nanovesicle, that encapsulate potentially therapeutic macromolecules such as proteins and nucleotides and deliver them across the BBB.^58,59^

Our SEs are synthesized in a microfluidic reactor, which allows tunability, control of size, charge, and encapsulation efficiency. SEs with encapsulated cargo form when the lipid and aqueous (carrying cargo) streams mix in the reactor.^58,60^ The lipids and other non-cargo components are Generally Regarded as Safe (GRAS) materials. Other similar LNP variants are a commonly used vehicle to deliver CRISPR to organs via blood administration,^32^ as well as other drugs like the COVID vaccine.^61^ As we have previously reported, SEs are similar in size to natural exosomes (<150 nm in diameter) and deformable.^58^ SEs are hypothesized to cross the BBB by uniquely deforming and squeezing through the tight junctions of vessels that form the BBB.^58^ SEs are similar to the lipid nanoparticles that have been used to deliver CRISPR for peripheral genetic diseases in humans.^56,57^

In this report, we show that SEs loaded with E4-specific CRISPR gRNA and CBE mRNA, administered by intravenous tail vein (IV) injection into E4 transgenic mice leads to editing E4 to E3 in brain and other organs, providing the first demonstration of the IV application of CRISPR to gene editing in brain.

## Materials and Methods

### Design and synthesis of gRNA

The key features of CRISPR CBE are gRNA and Cas9-linked cytosine deaminase. CBE requires exact PAM site spacing to the target base to allow the Cas9-linked deaminase to be positioned precisely over the target DNA site.^41^ The PAM site sequence must match the PAM specificity of the Cas9 being used, which for ApoE4 was set as ‘TG’ dinucleotide sequence. Using Benchling (https://benchling.com/pub/liu-base-editor), we analyzed the sequence around the E4 target ’C’ and identified two candidate PAM sites (both ‘TG’) in the E4 sequence at appropriate distances downstream from the E4 target base cytosine ’C’ (**Fig. 1A**). Relative to PAM, the deaminase peak activity window is typically from bases 4 to 8 in the gRNA sequence.

**Fig. 1.**
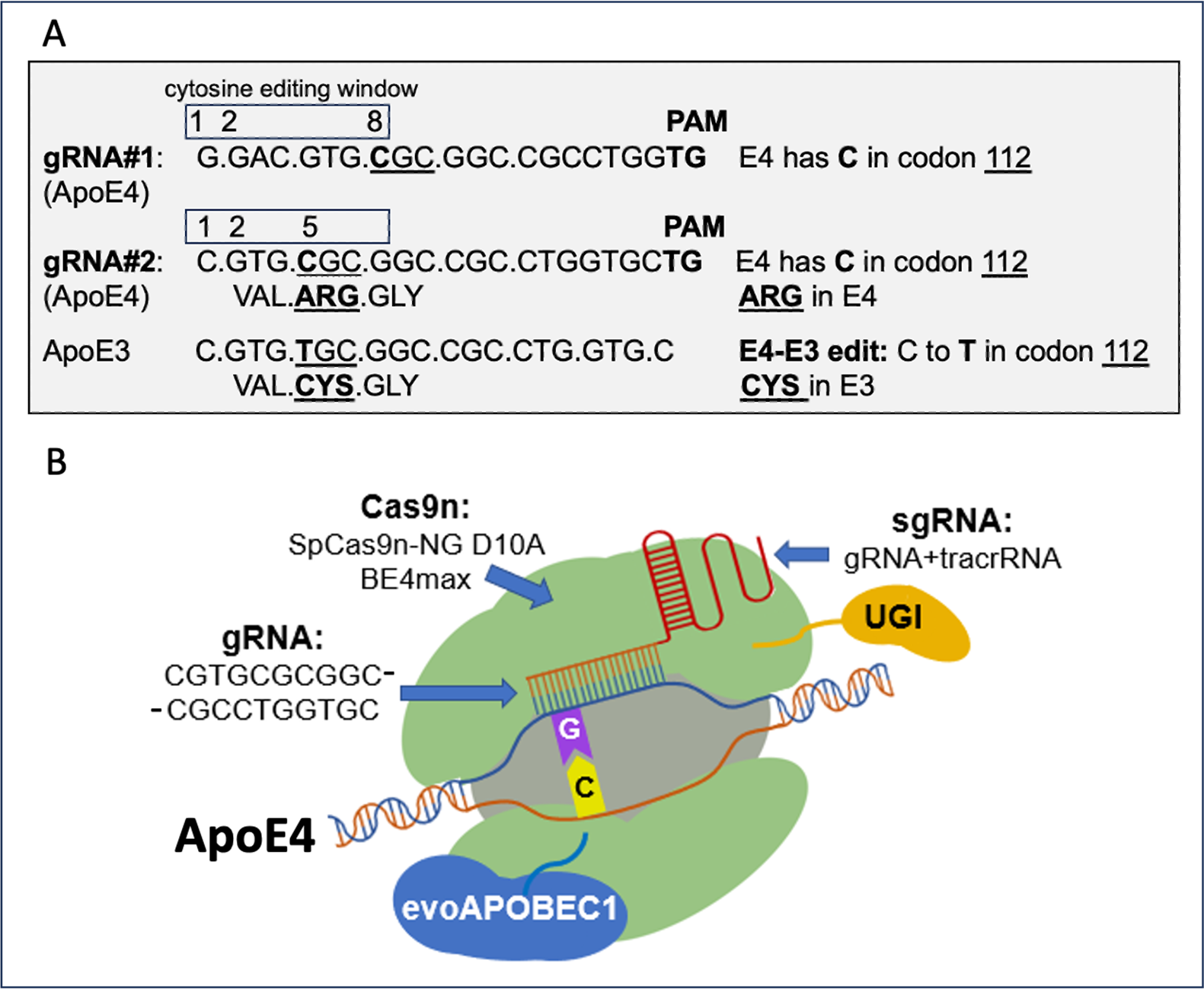
ApoE4 *gRNA variants, PAM site, and base editing of E4 ARG to E3 CYS*. (A) Shown are gRNAs #1 and #2 that target the *APOE4* sequence ‘**C**’ and align it over the cytosine deaminase cytosine editing window at positions #8 and #5, respectively (boxed). A ‘**C**’ to ‘**T**’ base edit in codon 112 results in an ARG to CYS substitution, creating ApoE3. (B) The CBE (Cas9n, evoAPOBEC1, and UGI), E4-specific gRNA and sgRNA, and ApoE4 gene complex are shown. The target cytosine ’**C**’ is shown near the cytosine deaminase evoAPOBEC1 enzyme.

Two ApoE4 gRNAs variants were designed that differ by the sequence of PAM sites they abut, thereby aligning the CBE cytosine deaminase activity window over slightly different positions over the target E4 ’C’ base. The first (gRNA#1) is the 20nt sequence abutting a PAM site ’TG’ which aligns the E4 cytosine ’C’ at position #8 in the CBE peak activity window; the second gRNA, gRNA#2, begins 3 bases downstream and extends 20nt to another ’TG’ PAM site, aligns the E4 ’C’ at position #5 in the CBE peak activity window.

The two gRNAs are linked with an 80 nucleotide (nt) trans-activating CRISPR (tracr)RNA that specifically binds Cas9 protein (**Fig. 1B**), and with appropriate base modifications (2’-O-methyl analogs, 3’-phosphorothionate linkages) for *in vivo* stability, were synthesized by Synthego (Redwood City, CA) at 150 nmol scale, purified, and lyophilized.

### CBE design and mRNA synthesis

The CBE is a fusion of 3 proteins: Cas9n - specifically Cas9n nickase D10A BE4max with ’NG’ sequence PAM specificity, cytosine deaminase evoAPOBEc1, and uracil glycosylase inhibitor (UGI). The complex of CBE, gRNA and sgRNA with the ApoE4 gene DNA sequence results in targeted base editing (**Fig. 1B**).

Two candidate CBEs, pBT375 and pYE1BE4max,^34^ were selected for synthesis and testing, that differ in several mutation-designed activities based on their targeting of PAM site sequence “TG”. Both plasmid expression vectors (ADDGENE) were modified with 3’ and 5’ UTR sequences (plasmids constructed by GeneScript, Piscataway, NJ) generated based on a protocol for CBE plasmid 3’ and 5’ UTR modifications. ^62^ The two CBE mRNAs were synthesized by Trilink (San Diego, CA) using *in vitro* transcription at 1 mg scale from PCR amplicons of the two expression plasmids.

### In vitro efficacy testing CRISPR gRNA and CBE mRNA variants in E4-N2A

The relative efficiency of editing E4 to E3 for each the four CRISPR gRNA+CBE mRNA combinations of two possible gRNA (#1 and #2) and two possible CBE mRNAs (pBT375 and pYE1BE4max) was tested in vitro in N2A cells that stably express an ApoE4 transgene (E4-N2A cells).^63^ CRISPR gRNA and CBE mRNA combinations were delivered to cells using Lipofectamine Messenger Max (LFMM), and after 2 days, the ApoE gene in genomic DNA was PCR amplified, and subjected to Sangar sequencing to identify CRISPR gRNA+CBE mRNA combinations with optimal E4 to E3 editing (**Supplementary Table S1).**

E4-N2A cells were cultured in DMEM medium and 10% FBS until 80% confluent, trypsinized, and transfected with the RNA combinations using Lipofectamine MessengerMax (LFMM, ThermoFisher) according to the manufacturer’s optimized protocol, modified using an protocol that improves transfection efficiency.^64^ Cells were trypsinized, resuspended in a small volume of serum-free DMEM to which the LFMM with gRNA+CBE mRNA (250 ng total RNA) was added, incubated 25°C, for 20 min, then plated at 25% confluence in 24-well plates and grown for 2 days. Cells were washed with PBS and genomic DNA extracted with Quick Extract (Lucigen), PCR amplified in the human ApoE gene exon 4 containing the E4 target base (see below), and the amplicons were Sanger sequenced (Laragen), and Sangar sequencing traces were quantified using EditR, as follows.

### Quantification of Sanger sequencing using EditR

Sanger sequencing data files (.ab1) and gRNA sequence were used to quantify the base traces corresponding to the E4 and E3 bases, using EditR (https://moriaritylab.shinyapps.io/editr_v10/), an open access online tool which generates a plot displaying base percentages (or editing efficiencies) at each base;^65^ it reduces the background signals and generates base/editing percentages based on a p-value of .05 with a detection limit for base editing of ∼7%; the statistical method in EditR assigns a probability that each background peak in the gRNA region is from noise – in other a words, a p-value for it being a product of base editing as opposed to noise.^65^

### PCR primers used for amplification of human ApoE gene exon 4

Primers were designed to amplify the human ApoE gene within exon 4:

Primer 1: GGCGCTGATGGACGAGAC

Primer 2: GTACACTGCCAGGCGCTTC

PCR amplification used a reaction containing Acuprime Taq polymerase (ThermoFisher) (which has the highest accuracy of Taq polymerases commercially available), the supplied buffer for amplifying genomic DNA, the supplied ’GC’ enhancer (since the region is ’GC’-rich), and thermocycled according the Acuprime manufacturer’s protocol, with annealing at 56°C and extension at 68°C. The PCR amplicon was 304 bp long, and the E3/E4 SNP site is at 119 bp from the P1 primer end.

### Microfluidic reactor synthesis of SEs

SEs with encapsulated cargo form when the lipid and aqueous (carrying cargo) streams mix in the microfluidic reactor.^58,60^ The lipids and other non-cargo components are Generally Regarded as Safe (GRAS) materials. The microfluidic reactor flow rates can be finely controlled to yield SEs with specific size (60<φ<500 nm), zeta potential (-50<ξ<50), and deformability. Once optimized, microfluidic synthesis of SEs is readily scalable to obtain larger amounts with good batch-to-batch reproducibility.

SE-CRISPR were synthesized in a Syrris microfluidic reactor by encapsulating 0.8 mg each of active gRNA and CBE mRNA or buffer (as a negative control). Total RNA (1.6 mg; 1:1 ratio of gRNA to CBE mRNA by mass; 60:1 ratio by molecule) was prepared in 4.8 mL of 10 mM NaCitrate, pH 4.0. The SE lipid mixture of DMPC (1,2-dimyristoyl-sn-glycero-3-phosphocholine): DPC (dihexadecyl phosphate, a negatively-charged lipid): CH (cholesterol) (catalog #s: Avantis Polar Lipid 850345P, Sigma Aldrich 2197-63-9, and Avantis Polar Lipid 700100P, respectively) at molar ratio of 4:2:2 was prepared by dissolving the lipids to a final concentration of 10 mg/mL in ethanol containing 2.3 % Span 80 w/v (Millipore Sigma cat# S6760). The lipid:RNA volume ratio was 3:1. After collection, the samples were dialyzed using 300 KDa MWCO dialysis tubing in sterile water at 4°C overnight. Samples were then aliquoted, lyophilized, and stored at -20°C. SE-empty was similarly synthesized in the microfluidic reactor as above absent the CRISPR RNA components.

### SE quality control assays

SE vesicle size was determined by Dynamic Light Scattering (DLS; DynaPro Plate Reader III, Wyatt Technologies) and SE concentration was determined by Nanotracker using a vesicle-specific fluorescent dye (exoGlo; AlphaNanoTech). RNA encapsulation efficiency was measured using the Ribogreen assay (Thermo Fischer) in an aliquot of lyophilized SE-CRISPR that had been resuspended in PBS and lysed using 5% Triton X100. RNA encapsulation efficiency was calculated using: ‘lysed RNA (total RNA) - unlysed RNA’ (unincorporated RNA) / total RNA). RNA integrity was determined by gently purifying RNA from a post-lyophilized aliquot using a Qiagen RNEasy kit with RLTplus buffer (high detergent lysis), followed by analysis using Tapestation.

### In vitro efficacy testing of SE-CRISPR in human Kelly neuroblastoma cells

The Kelly human neuroblastoma cell line has an endogenous ApoE genotype of E3/E4. ^66^ To assess the ability of SEs which encapsulate the CRISPR RNA combination with the highest editing efficiency shown in E4-N2A cells (gRNA #2 + pBT375 mRNA) to edit the ApoE4 gene, negatively-charged lipid SEs were synthesized to encapsulate CRISPR RNAs gRNA#2+CBE mRNA pBT375, and used to treat the cells.

Kelly cells were plated to 15% confluence in 24-well plates, and treated once with either SE-CRISPR or SE-empty, grown for 5 days, cells were harvested, and prepared like the E4-N2A cells above, for PCR amplicon Sangar sequencing and quantification using EditR. The same CRISPR RNAs were also delivered to Kelly cells using Lipofectamine Messenger Max, as described above for E4-N2A cells.

### In vivo testing of SE-CRISPR in E4-5XFAD mice

The ApoE4-targeted replacement-5XFAD (E4-5XFAD) mice used for in vivo testing of SE-CRISPR express human ApoE4 from a targeted-replacement ApoE4 gene and are homozygous for ApoE4.^67^ The mice also express human amyloid precursor protein (APP) with K670N/M671L, I716V, and V717I mutations, as well as presenilin 1 (PS1) with M146L and L286V mutations under the control of the neuron-specific mouse thymocyte differentiation antigen 1 theta (Thy-1) promotor.^67^

An aliquot of lyophilized SE-CRISPR was resuspended in PBS at a concentration of 0.15 mg total RNA/100 μL, which is a dose of 5 mg RNA/kg mouse weight (previous reported dose range for liver targeting is between 1-5 mg/kg mouse ^68,69^); the suspension was filtered at 0.2 μm, producing a clear solution. SEs containing only buffer (SE-empty) were used as controls. A volume of 100 μL SE-CRISPR (or SE-empty) was injected IV into the tail vein of an E4-5XFAD mouse (n = 1; 6 month old). Five (5) days post-injection, mice were deeply anesthetized, blood was collected transcardially, and mice underwent extensive perfusion with 37°C PBS until the liver had become pale plus an additional 2 minutes of perfusion. The day 5 euthanasia time point was chosen based on published reports,^70,68^

Brain and liver tissue were collected and buffy coat (comprising white blood cells, WBCs) were isolated from blood. Genomic DNA was prepared using Quick Extract (Lucigen) and column purified. The ApoE4 region was PCR amplified, amplicons purified, followed by quality control for yield and purity, and used for library construction and Illumina Short-Read deep sequencing (performed the UCLA TCGB Core), to a depth of 27+/- 6 million reads per sample. NGS data was analyzed for the number of E4 and E3 sequence molecules. The %E3 was calculated as: #E3 counts / (#E4 counts + #E3 counts) x 100%.

Bystander editing, which results from CRISPR editing the cytosine near the E4-specific cytosine, was also assessed (see **Fig. 4A**).

The scheme for *in vivo* testing of SE-CRISPR in ApoE4 targeted-replacement (E4-5XFAD) mice is shown in **Fig. 2**, and comprises SE-CRISPR synthesis, dialysis, lyophilization, storage, re-suspension just before use, IV injection, 5-day incubation, euthanasia, tissue collection, and analysis.

**Fig. 2.**
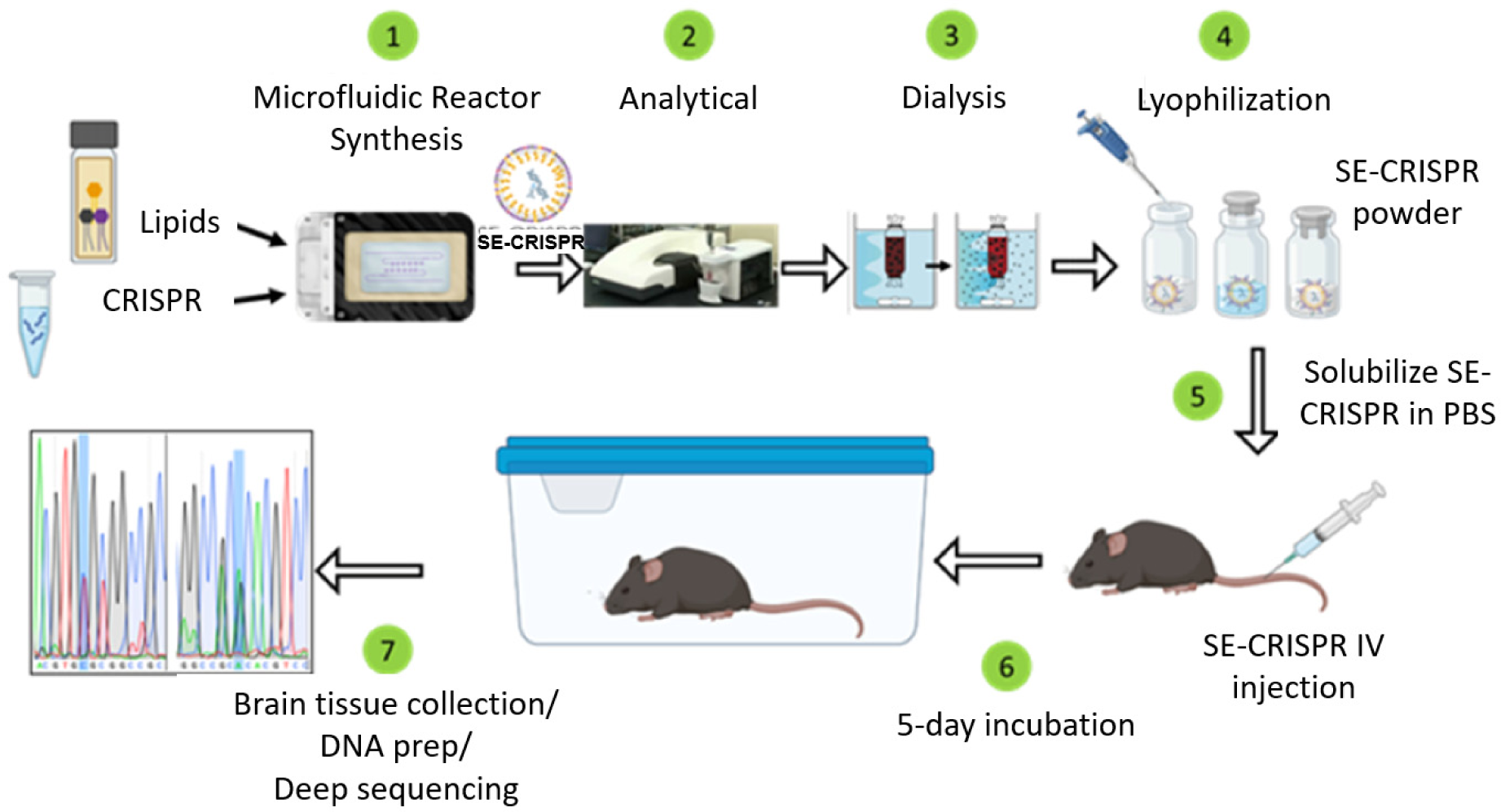
Scheme for in vivo testing of SE-CRISPR in mice. The experimental scheme for in vivo testing of SE-CRISPR in mice comprises the steps as shown.

### Determination of E3 mRNA levels

Successful editing of the E4 gene to E3 is expected to result in functional gene expression and the production of E3 mRNA in the tissues from the SE-CRISPR-treated mouse. For assay of E3 mRNA, RNA was isolated from brain tissue of the SE-CRISPR-treated mouse and control mouse (the same mice used to measure %E3 DNA editing), converted to cDNA, PCR amplified, sequencing library constructed, and sequenced by NGS. E3 mRNA counts were converted to %E3 mRNA = E3 mRNA counts / (E3 counts + E4 counts), and compared to the %E3 gene editing.

### Brain Capillary Depletion

Brain tissue was fractioned into capillary-enriched and parenchymal (capillary-depleted) fractions using the Tenbroek homogenization and dextran centrifugation method.^71^ Briefly, one brain hemisphere was homogenized using a Tenbroek homogenizer (Corning) in 18% dextran 70,000 (Millipore) and centrifuged at 3260xg for 40 minutes at 4°C. The supernatant was carefully removed and the pellet was resuspended in HBSS. Aliquots were taken for assay of acid phosphatase activity, which is preferentially expressed by capillary endothelial cells, and for measuring E3 editing.

## Results

### Identification of gRNA and CBE RNA combination with highest editing efficiency

Of the four combinations of gRNAs that target the E4 cytosine and CBE mRNA (**Supplementary Table S1**), the combination of gRNA #2 and pBT375 CBE mRNA exhibited the highest editing efficiency of ApoE4 genomic DNA in E4-N2A cells, as detected by Sanger sequencing. The findings from Sanger sequencing of E4-N2A cells transfected with combination gRNA#2 + pBT375 CBE mRNA are shown in **Fig. 3**. The DNA strand containing the target base ’**C**’ in ApoE4 is highlighted blue (**Fig. 3A** and **Fig 3B**)(’**C**’ is shown in blue traces). Results of editing (**Fig. 3B**) show red ’**T**’ (shown in red traces) is superimposed on the blue ’**C**’ trace (labelled as **’C/T’**), and they are equally high. These traces were quantified for editing efficiency (relative abundance of ’**C**’ and ’**T**’ at the target base site) using EditR ^65^ which showed 51% edited ’**T**’. Similarly and to validate these results on one DNA strand (**Fig. 3B**), traces from the complimentary DNA strand are shown in **Fig. 3C** and **Fig. 3D**, where the control cell E4 target position ’**G’** (**Fig. 3C**) is edited to ’**A’** (labelled as ’**G/A’**) (**Fig. 3D**). EditR analysis showed 54% edited ’**A’**. Therefore, analysis of both DNA strand’s editing indicate the combination of gRNA#2 + pBT375 CBE mRNA was very efficient, relative to ∼50% transfection efficiency.^64^ In contrast, the other three combinations of gRNA+CBE mRNA had lower editing efficiency of <15% (**Supplementary Table S1**).

**Fig. 3.**
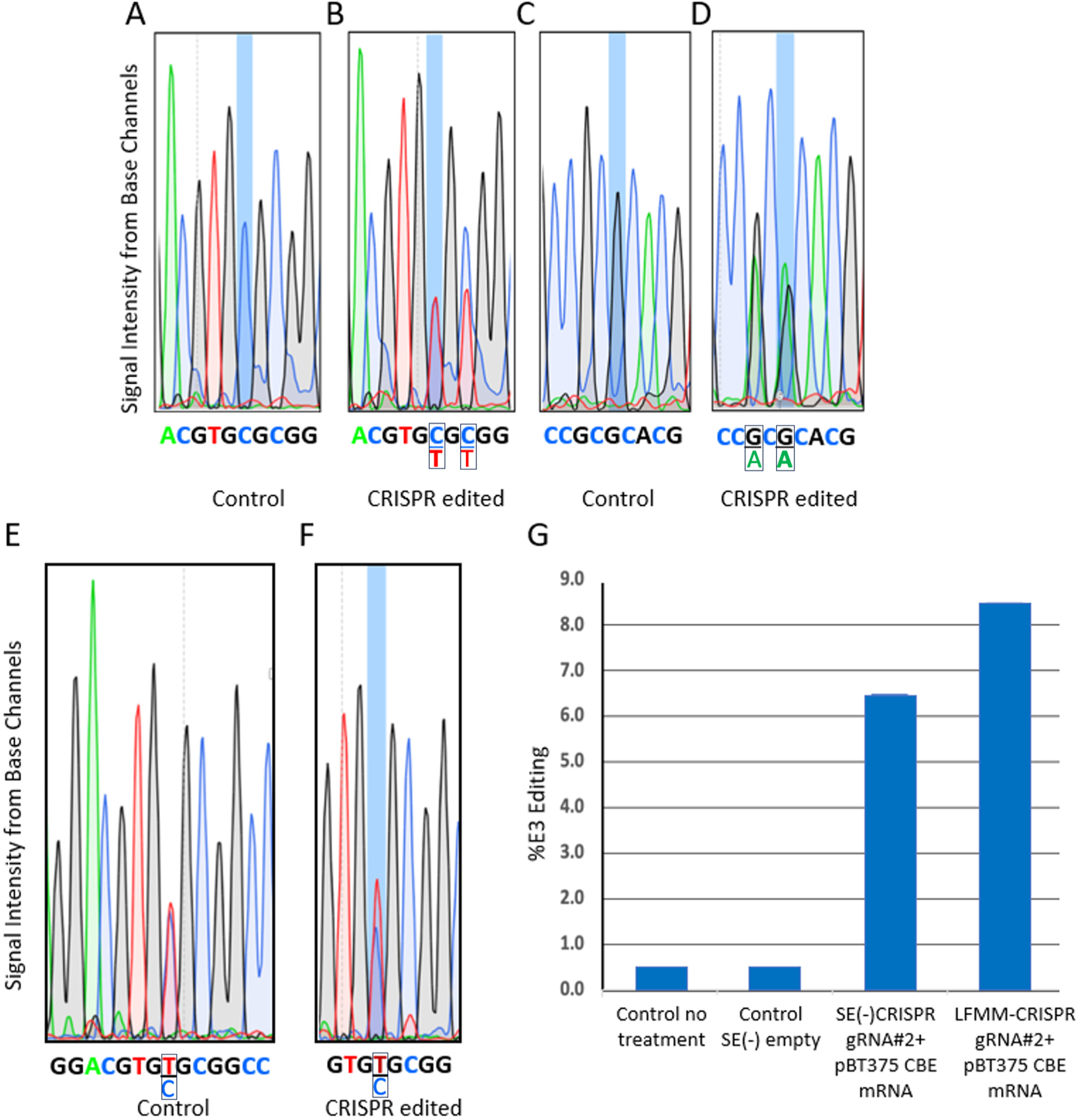
Sanger sequencing traces from CRISPR-edited and control E4-N2A and Kelly neuroblastoma cells. DNA strand base traces from Sanger sequencing of E4 PCR amplicons from E4-N2A cells are shown for (A) target ’**C**’ (highlighted blue) control cells; (B) target ’**C**’ (highlighted blue) and edited ’**T**’ from cells transfected with gRNA(#2) + pBT375 CBE mRNA (box labelled **’C/T**’); (C) the complimentary DNA strand target ’**G**’ (highlighted blue) from control cells; and (D) target ’**G**’ (highlighted blue) and edited ’**A**’ from cells transfected with gRNA(#2) + pBT375 CBE mRNA (box labelled ’**G/A**’). Bystander editing traces are shown in (B) and (D), labelled as boxed ’C/T’ and ’G/A’ at the position 2 bases from the target ’**C**’ or ’**G**’ base. Traces are similarly shown for human Kelly neuroblastoma from (E) control and (F) CRISPR edited cells. (G) The %E3 base editing as quantified by EditR, calculated as %’T’ treated-%’T’ untreated, is shown for no treatment, SE-empty, SE-CRISPR, and lipofectamine messenger max (LFMM)-CRISPR in Kelly cells. For A-F, the chromatograms represent the migration of labeled sequencing products via capillary electrophoresis. The bases represented by each of the four color channels are: green: ’A’; blue: ’C’; black: ’G’; red: ’T’. Edited E4 bases are highlighted in blue. Note that in (B), for the target ’**C/T’** trace, the red ’**T**’ trace is coincident with the blue **’C’** trace.

Regarding bystander editing, the bystander ’C’ located two bases 3’ of the E4 target ’**C**’ (**Fig. 3A**), is also substantially edited to ’T’ (**Fig. 3B**); likewise, on the complimentary strand, the ’G’ (Fig .3C) is edited to ’A’ (**Fig. 3D**)). This bystander editing is expected within the CBE editing window (**Fig. 1A**).^72^ The resultant ’TGT’ codon would still result in the E3 cysteine.

### Characterization of SE-CRISPR

The average SE-CRISPR particle size, as determined by DLS and Nanotracker (NTA) analysis, was ∼90 nm (**Supplementary Fig. S1**). NTA detected 26% of total particles were vesicles, and a vesicle concentration of 2.25 X 10^9^ vesicles/mL in an aliquot from the preparation used for injection. This resulted in injection of 2.25 X 10^8^ vesicles/100uL dose.

RNA encapsulation efficiency, measured using the Ribogreen assay, showed 93% encapsulation of the gRNA + pBT375 CBE mRNA. RNA integrity, analyzed using Tapestation, showed intact peaks for the 6000nt mRNA and the 100nt gRNA (sgRNA is 20nt gRNA+80nt TracrRNA (**Supplementary Fig. S2**).

### SE-CRISPR E4 to E3 editing in human ‘Kelly’ neuroblastoma cells

The ability of SEs to deliver the CRISPR RNAs gRNA#2 + pBT375 CBE mRNA was evaluated in vitro in Kelly neuroblastoma cells that has ApoE genotype E3/E4.^66^ The Sanger sequencing traces from genomic DNA of Kelly cells transfected with SE-CRISPR (gRNA#2+pBT375 CBE mRNA) and then culture for 5 days is shown in **Fig. 3**.

Kelly cells treated with SE-empty (**Fig. 3E**) display the E3/E4 heterozygosity of the Kelly cell line - ’C’ and ’T bases at 50% each at the target base (overlapping red ’T’ and blue ’C’ trace; labelled as ’T/C’). SE-CRISPR-treated cells (**Fig. 3F**) show increased ’T’ base (red trace). Quantification of the edited percent of ’T’ (calculated as %’T’ treated-%’T’ untreated) indicated 6.5% editing of E4 to E3 by SE-CRISPR, similar to that achieved by delivering the same CRISPR RNAs using Lipofectamine Messenger Max (8.5% editing) (**Fig. 3G**).

### SE-CRISPR E4 to E3 editing in brain, liver and white blood cells of E4-5XFAD mice

E4-5XFAD mice were injected IV with SE-CRISPR or SE-empty and after 5 days, tissues were analyzed by NGS for editing the E4 gene to E3. As shown in **Fig. 4A**, NGS analysis of tissue collected from the mouse treated with SE-CRISPR gRNA#2 + pBT375 CBE mRNA revealed that SE-CRISPR successfully edited E4 to E3 in liver (0.81 %E3), brain (0.14 %E3), and buffy coat/WBCs (0.36 %E3). %E3 was calculated as: #E3 counts / (#E4 counts + #E3 counts) x 100%. Data are presented in **Supplementary Table S2.**

**Fig. 4.**
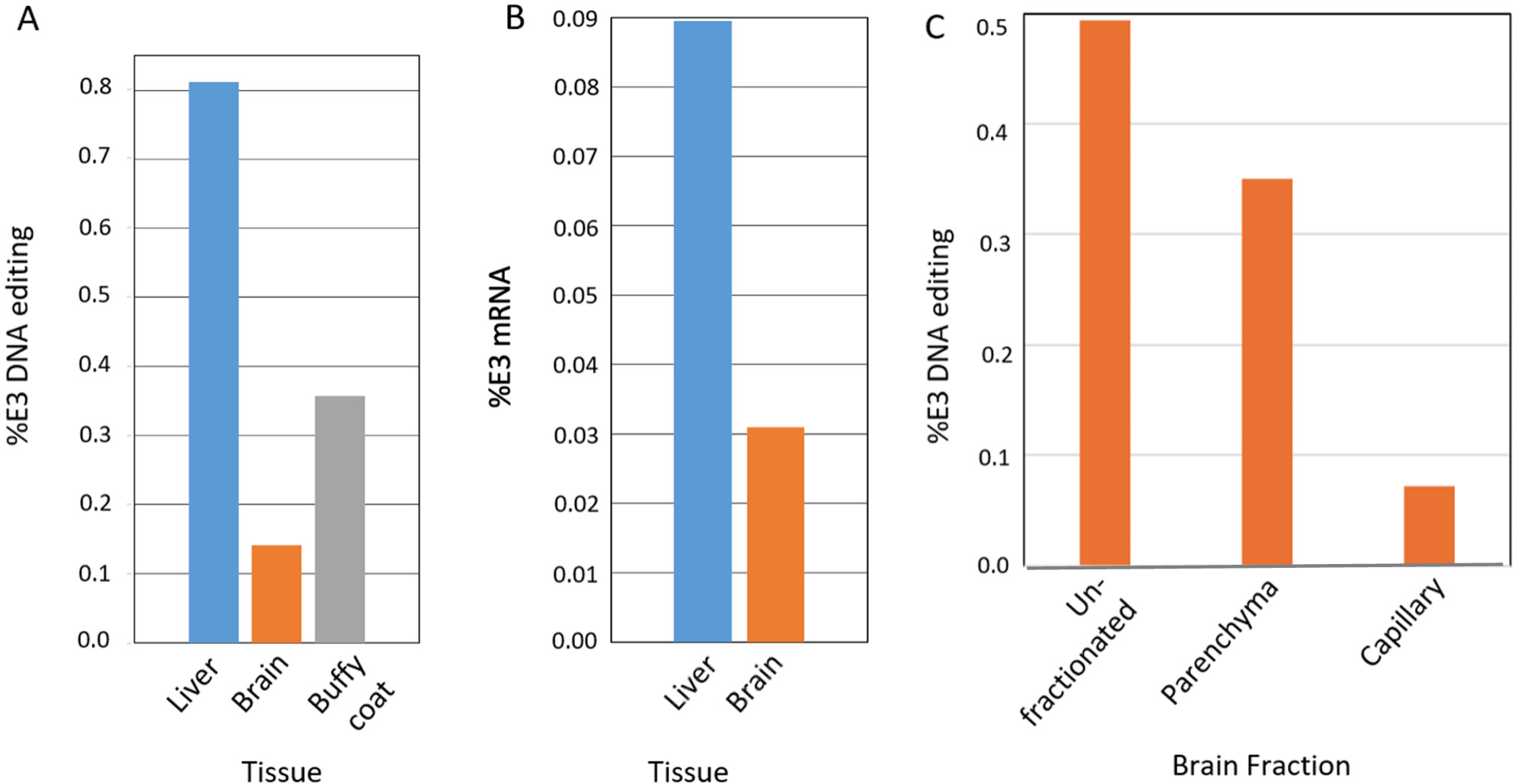
SE-CRISPR editing of E4 to E3 in different tissues and brain fractions of an ApoE4-5XFAD mouse. (A) The %E3 DNA editing (normalized to SE-empty) is shown for brain, liver, and buffy coat (white blood cells) 5 days after IV injection of SE-CRISPR into E4-5XFAD mice. (B) The %E3 mRNA normalized to SE-empty, in liver and brain, are shown. **Supplementary Table S2** presents the values for %E3 DNA editing (n = 1). CRISPR = gRNA#2+ pBT375 CBE mRNA. (C) Editing %E3 measured by NGS is shown for unfractionated brain and for brain fractions enriched for parenchyma and capillary cells, from the brain of an E4 mouse treated with SE-CRISPR. E3 counts were normalized to the total NGS reads per sample and to the reads obtained from the SE-empty-treated mouse.

Bystander editing was observed in all three tissues at levels < 0.025%, consistent with the expected lower level of editing compared to that of the targeted E4 to E3 editing.^72^

To evaluate the functional outcome of editing the E4 gene to E3, mRNA in the brain and liver of SE-CRISPR-treated mice were analyzed by NGS for the level of E3 mRNA. SE-CRISPR-treated mice had 0.09% E3 mRNA in liver and 0.03% in brain (**Fig. 4B**).

### SE-CRISPR E4 to E3 editing in brain of E4-5XFAD mice after capillary depletion

To evaluate whether E3 editing by SE-CRISPR was in brain capillary endothelial cells and/or brain parenchymal cells (neurons and glia), one brain hemisphere from the SE-CRISPR-treated mouse and one from the SE-empty-treated mouse were fractionated into capillary-enriched and capillary-depleted fractions. In this method, the capillaries are pelleted and the supernatant is depleted of capillaries, and represents brain parenchymal-enriched cells.^71^ Using this method we generated enriched fractions of capillary and parenchymal cells, based on enrichment of acid phosphatase in the capillary fraction (**Supplementary Fig. S3**).^71^

E4 to E3 editing in unfractionated, parenchymal, and capillary brain fractions by NGS is shown in **Fig. 4**. Compared to that in the unfractionated brain, overall gene editing in the parenchymal fraction was 71%, and in the capillary fraction 14%, indicating that the E3 editing in unfractionated brain was largely accounted for by E3 editing in the parenchymal-enriched fraction (**Supplementary Table S3**). The loss of some sample in the fractionation process could likely account for the %E3 editing not evident in either the parenchymal or capillary samples.

## Discussion

We successfully designed, tested, and identified an optimized CRISPR gRNA+CBE combination that edits the E4 gene to E3 in vitro in murine neuroblastoma cells that express ApoE4 (E4-N2a). We then encapsulated the gRNA+CBE combination with the highest E4 editing in vitro in SE particles, and used them to treat Kelly human neuroblastoma cells, and demonstrated successful editing. Further, in a proof-of-concept study in vivo using the optimized CRISPR combination, a modest level of E4 to E3 editing (0.14%) after single IV dose of SE-CRISPR was demonstrated in brain tissue of an ApoE4-expressing AD model mouse.

E4 to E3 editing was also observed in in liver (0.8 %E3) and WBCs (0.36 %E3). Editing in WBCs was anticipated because of the IV delivery route, which would enable the SE-CRISPR to be in direct contact with WBCs. Editing in liver was also expected because numerous studies have shown that LNP-delivered CRISPR edits liver cells because of their robust uptake of blood-borne LNPs.^73^

Bystander editing was very low, < 0.025%, in brain, liver and WBCs. Because the bystander editing-generated sequence codon ’TGT’ does not exist in nature, it provides evidence that the ’TGT’ sequence, as well as the ’TGC’ E3 sequence (generated in E4 mice), has been created by CRISPR editing. When CRISPR edits both cytosines in the E4 arginine codon (’CGC’) to ’TGT’, the resulting codon remains an E3 cysteine (the third position ’T’ is redundant, a silent mutation). Further, the level of bystander editing (when calculated by %bystander/%E3) was consistent across the three tissues examined, where bystander editing was 1.5%-2.4% that of E3 editing.

For brain, the %E3 of 0.14% represents editing of about 1 in ∼700 cells, or for the mouse brain with ∼3.2 x 10^7^ total cells ^74^, ∼4.3 x 10^6^ E3 cells (assuming heterozygous editing per cell). For liver, the %E3 of 0.8% represents editing of about 1 in ∼125 cells. These calculations can be used to make a simplistic prediction about how many vesicles need to be injected to achieve a target of 50% editing E4 to E3 in brain; this 50% editing target would represent the gene dosage of an E3/E4 heterozygote which should partially neutralize E4/E4-specific AD-relevant phenotypes. Extrapolating from the mouse injected with 2.25 x 10^6^ SE-CRISPR vesicles that achieved 0.14 %E3 in brain, to achieve 50% E3 editing in one single injection, it is predicted that the number of vesicles needed would be 8 x 10^10^ (80 billion). In future studies, this could be achieved with a series of injections over time.

The levels of E3 mRNA in SE-CRISPR-treated mice was 0.09% E3 mRNA in liver and 0.03% in brain, which compared to the level of E3 editing in the same tissues, are between 11%-22% the level of E3 gene DNA. This is consistent with E3 mRNA expression not yet achieving equilibrium by 5 days post IV; with additional time, it is expected the pre-existing E4 mRNA would be turned over, and that the levels of E3 and E4 mRNA would reflect their gene ratios.

We confirmed that E4 to E3 editing in the brain was largely parenchymal, with a smaller contribution in unfractionated brain tissue due to editing in capillary endothelial cells. Editing in the parenchyma is evidence that the SE-CRISPR has transcytosed the BBB into brain parenchyma.^70,71^ This is consistent with the ability of SE particles to cross the BBB, hypothesized to be attributed to their lipid-based deformability.^58^ This is a critical demonstration in the development of SE-CRISPR as a translatable drug. This also demonstrates the potential of SEs as a drug-delivery platform that has the ability to deliver cargo to the brain that might not otherwise pass the BBB.

Editing the E4 gene to E3, and the resultant ApoE3 protein production, is predicted to not only affect the edited neuronal and glial cells but neighboring cells as well, because ApoE produced by glial cells in the brain is secreted and then taken up by other cells via receptor-mediated endocytosis.^19^

Herein we have demonstrated initial proof-of-concept for editing ApoE4 to E3 in the brain after a single IV infusion of SE-CRISPR. Future studies will focus on increasing the editing percent of the current SE-CRISPR candidate by increased SE particles in the dosing, and brain penetrance through surface modifications that increase blood half-life. We plan to assess relationships between E3 protein expression in the brain and SE-CRISPR dose/exposure to therapeutic modulation E4-specific phenotypes over time. Based on the number of cells in mouse brain showing E4 to E3 editing after a single IV infusion, the translation of this therapy for clinical testing is feasible. Successful development of an SE-CRISPR IV infusion therapy could potentially be beneficial to millions of AD and Mild Cognitive Impairment (MCI) ^75,76^ patients having one or two copies of ApoE4,^5^ as a monotherapy and/or as part of multi-modal therapy with approved anti-amyloid antibodies such as aducanumab and lecanemab.

## Author contributions

BT designed, supervised or carried out experiments, prepared data for analysis, and co-wrote the manuscript; JC co-developed the SE platform, synthesized SE particles, and performed DLS quality control; CZ performed tail vein injection and animal perfusion and sacrifice for *in vivo* studies; DBK advised on experimental design and data interpretation; GEM assisted with gRNA and CBE design and plasmid 3’ and 5’ UTR modifications; PS contributed to experimental design, data interpretation, and co-wrote the manuscript; VJ conceived the project, co-designed experiments, interpreted data, and edited the manuscript. All authors approved the manuscript.

## Acknowledgements

We thank Whitaker Cohn, Ph.D. for technical assistance, Sahiba Beniwal for helping with mouse colony maintenance, and Alicia Li and Kierra Zaneskifor helping on the microfluidic synthesis of SEs.

## References

1 Belloy, M. E., Napolioni, V. & Greicius, M. D. A Quarter Century of APOE and Alzheimer’s Disease: Progress to Date and the Path Forward. Neuron 101, 820–838, doi:10.1016/j.neuron.2019.01.056 (2019).

2 Corder, E. H. et al. Gene dose of apolipoprotein E type 4 allele and the risk of Alzheimers disease in late onset families. Science 261, 921, doi:10.1126/science.8346443 (1993).

3 Farrer, L. A. et al. Apolipoprotein E genotype in patients with Alzheimer’s disease: implications for the risk of dementia among relatives. Annals of neurology 38, 797–808, doi:10.1002/ana.410380515 (1995).

4 Genin, E. et al. APOE and Alzheimer disease: a major gene with semi-dominant inheritance. Molecular psychiatry 16, 903–907, doi:10.1038/mp.2011.52 (2011).

5 Pires, M. & Rego, A. C. Apoe4 and Alzheimer’s Disease Pathogenesis-Mitochondrial Deregulation and Targeted Therapeutic Strategies. Int J Mol Sci 24, doi:10.3390/ijms24010778 (2023).

6 Reiman, E. M. et al. Exceptionally low likelihood of Alzheimer’s dementia in APOE2 homozygotes from a 5,000-person neuropathological study. Nature communications 11, 667, doi:10.1038/s41467-019-14279-8 (2020).

7 Koriath, C. et al. ApoE4 lowers age at onset in patients with frontotemporal dementia and tauopathy independent of amyloid-β copathology. Alzheimer’s & dementia (Amsterdam, Netherlands) 11, 277–280, doi:10.1016/j.dadm.2019.01.010 (2019).

8 Shi, Y. et al. ApoE4 markedly exacerbates tau-mediated neurodegeneration in a mouse model of tauopathy. Nature 549, 523–527, doi:10.1038/nature24016 (2017).

9 Neitzel, J. et al. ApoE4 associated with higher tau accumulation independent of amyloid burden. Alzheimer’s & Dementia 16, e046206, doi: 10.1002/alz.046206 (2020).

10 Baek, M. S. et al. Effect of APOE ε4 genotype on amyloid-β and tau accumulation in Alzheimer’s disease. Alzheimers Res Ther 12, 140, doi:10.1186/s13195-020-00710-6 (2020).

11 Braak, H. & Braak, E. Neuropathological stageing of Alzheimer-related changes. Acta Neuropathol 82, 239–259, doi:10.1007/bf00308809 (1991).

12 Hardy, J. A. & Higgins, G. A. Alzheimer’s disease: the amyloid cascade hypothesis. Science 256, 184–185, doi:10.1126/science.1566067 (1992).

13 Bancher, C. et al. Accumulation of abnormally phosphorylated τ precedes the formation of neurofibrillary tangles in Alzheimer’s disease. Brain Research 477, 90–99, doi: 10.1016/0006-8993(89)91396-6 (1989).

14 Goedert, M. Tau protein and the neurofibrillary pathology of Alzheimer’s disease. Trends in neurosciences 16, 460–465 (1993).

15 Cao, J. et al. ApoE4-associated phospholipid dysregulation contributes to development of Tau hyper-phosphorylation after traumatic brain injury. Sci Rep 7, 11372, doi:10.1038/s41598-017-11654-7 (2017).

16 Liu, C. C. et al. ApoE4 Accelerates Early Seeding of Amyloid Pathology. Neuron 96, 1024–1032.e1023, doi:10.1016/j.neuron.2017.11.013 (2017).

17 Mahley, R. W., Huang, Y. & Weisgraber, K. H. Detrimental effects of apolipoprotein E4: potential therapeutic targets in Alzheimer’s disease. Curr Alzheimer Res 4, 537–540 (2007).

18 Vassar, R. Seeds of Destruction: New Mechanistic Insights into the Role of Apolipoprotein E4 in Alzheimer’s Disease. Neuron 96, 953–955, doi:10.1016/j.neuron.2017.11.022 (2017).

19 Serrano-Pozo, A., Das, S. & Hyman, B. T. APOE and Alzheimer’s disease: advances in genetics, pathophysiology, and therapeutic approaches. The Lancet. Neurology 20, 68–80, doi:10.1016/s1474-4422(20)30412-9 (2021).

20 Johnson, J. K., McCleary, R., Oshita, M. H. & Cotman, C. W. Initiation and propagation stages of beta-amyloid are associated with distinctive apolipoprotein E, age, and gender profiles. Brain Res 798, 18–24, doi:10.1016/s0006-8993(98)00363-1 (1998).

21 Mahley, R. W. & Huang, Y. Alzheimer disease: multiple causes, multiple effects of apolipoprotein E4, and multiple therapeutic approaches. Annals of neurology 65, 623–625, doi:10.1002/ana.21736 (2009).

22 Vance, J. M. et al. Report of the APOE4 National Institute on Aging/Alzheimer Disease Sequencing Project Consortium Working Group: Reducing APOE4 in Carriers is a Therapeutic Goal for Alzheimer’s Disease. Annals of neurology, doi:10.1002/ana.26864 (2024).

23 Xiong, M. et al. APOE immunotherapy reduces cerebral amyloid angiopathy and amyloid plaques while improving cerebrovascular function. Sci Transl Med 13, doi:10.1126/scitranslmed.abd7522 (2021).

24 Kantor, B., Rittiner, J., Nicholls, P. J. & Chiba-Falek, O. APOE-targeted epigenome therapy for late onset Alzheimer’s disease. Alzheimer’s & Dementia 18, e060974, doi: 10.1002/alz.060974 (2022).

25 Offen, D., Rabinowitz, R., Michaelson, D. & Ben-Zur, T. Towards gene-editing treatment for alzheimer’s disease: ApoE4 allele-specific knockout using a CRISPR cas9 variant. Cytotherapy 20, S18, doi:10.1016/j.jcyt.2018.02.036 (2018).

26 Adji, A. S. et al. A Review of CRISPR Cas9 for Alzheimer’s Disease: Treatment Strategies and Could target APOE e4, APP, and PSEN-1 Gene using CRISPR cas9 Prevent the Patient from Alzheimer’s Disease? Open Access Macedonian Journal of Medical Sciences 10, 745–757, doi:10.3889/oamjms.2022.9053 (2022).

27 Nelson, M. R. et al. The APOE-R136S mutation protects against APOE4-driven Tau pathology, neurodegeneration and neuroinflammation. Nature Neuroscience 26, 2104–2121, doi:10.1038/s41593-023-01480-8 (2023).

28 Mahley, R. W. & Huang, Y. Small-molecule structure correctors target abnormal protein structure and function: structure corrector rescue of apolipoprotein E4-associated neuropathology. J Med Chem 55, 8997–9008, doi:10.1021/jm3008618 (2012).

29 Wang, C. et al. Gain of toxic apolipoprotein E4 effects in human iPSC-derived neurons is ameliorated by a small-molecule structure corrector. Nature Medicine 24, 647–657, doi:10.1038/s41591-018-0004-z (2018).

30 Sun, Y. Y., Wang, Z. & Huang, H. C. Roles of ApoE4 on the Pathogenesis in Alzheimer’s Disease and the Potential Therapeutic Approaches. Cell Mol Neurobiol 43, 3115–3136, doi:10.1007/s10571-023-01365-1 (2023).

31 Weisgraber, K. H. Apolipoprotein E distribution among human plasma lipoproteins: role of the cysteine-arginine interchange at residue 112. J Lipid Res 31, 1503–1511 (1990).

32 Porto, E. M., Komor, A. C., Slaymaker, I. M. & Yeo, G. W. Base editing: advances and therapeutic opportunities. Nature reviews. Drug discovery 19, 839–859, doi:10.1038/s41573-020-0084-6 (2020).

33 Evanoff, M. & Komor, A. C. Base Editors: Modular Tools for the Introduction of Point Mutations in Living Cells. Emerg Top Life Sci 3, 483–491, doi:10.1042/etls20190088 (2019).

34 Komor, A. C., Kim, Y. B., Packer, M. S., Zuris, J. A. & Liu, D. R. Programmable editing of a target base in genomic DNA without double-stranded DNA cleavage. Nature 533, 420–424, doi:10.1038/nature17946 (2016).

35 Rohn, T. T., Kim, N., Isho, N. F. & Mack, J. M. The Potential of CRISPR/Cas9 Gene Editing as a Treatment Strategy for Alzheimer’s Disease. Journal of Alzheimer’s disease & Parkinsonism 8, doi:10.4172/2161-0460.1000439 (2018).

36 Safieh, M., Korczyn, A. D. & Michaelson, D. M. ApoE4: an emerging therapeutic target for Alzheimer’s disease. BMC Medicine 17, 64, doi:10.1186/s12916-019-1299-4 (2019).

37 Wadhwani, A. R., Affaneh, A., Van Gulden, S. & Kessler, J. A. Neuronal apolipoprotein E4 increases cell death and phosphorylated tau release in alzheimer disease. Annals of neurology 85, 726–739, doi:10.1002/ana.25455 (2019).

38 Lin, Y. T. et al. APOE4 Causes Widespread Molecular and Cellular Alterations Associated with Alzheimer’s Disease Phenotypes in Human iPSC-Derived Brain Cell Types. Neuron 98, 1294, doi:10.1016/j.neuron.2018.06.011 (2018).

39 Lu, L., Yu, X., Cai, Y., Sun, M. & Yang, H. Application of CRISPR/Cas9 in Alzheimer’s Disease. Frontiers in neuroscience 15, 803894, doi:10.3389/fnins.2021.803894 (2021).

40 Zhao, J. et al. APOE4 exacerbates synapse loss and neurodegeneration in Alzheimer’s disease patient iPSC-derived cerebral organoids. Nature communications 11, 5540, doi:10.1038/s41467-020-19264-0 (2020).

41 Caso, F. & Davies, B. Base editing and prime editing in laboratory animals. Laboratory animals, 23677221993895, doi:10.1177/0023677221993895 (2021).

42 Vasquez, C. A., Cowan, Q. T. & Komor, A. C. Base Editing in Human Cells to Produce Single-Nucleotide-Variant Clonal Cell Lines. Current protocols in molecular biology 133, e129, doi:10.1002/cpmb.129 (2020).

43 Yeh, W. H., Chiang, H., Rees, H. A., Edge, A. S. B. & Liu, D. R. In vivo base editing of post-mitotic sensory cells. Nature communications 9, 2184, doi:10.1038/s41467-018-04580-3 (2018).

44 Porto, E. M. & Komor, A. C. In the business of base editors: Evolution from bench to bedside. PLoS Biol 21, e3002071, doi:10.1371/journal.pbio.3002071 (2023).

45 McAuley, G. E. et al. Human T&#xa0;cell generation is restored in CD3&#x3b4; severe combined immunodeficiency through adenine base editing. Cell 186, 1398–1416.e1323, doi:10.1016/j.cell.2023.02.027 (2023).

46 Hanafy, A. S., Schoch, S. & Lamprecht, A. CRISPR/Cas9 Delivery Potentials in Alzheimer’s Disease Management: A Mini Review. Pharmaceutics 12, doi:10.3390/pharmaceutics12090801 (2020).

47 Thompson, T. How CRISPR gene editing could help treat Alzheimer’s. Nature 625, 13–14, doi:10.1038/d41586-023-03931-5 (2024).

48 Bhardwaj, S. et al. CRISPR/Cas9 gene editing: New hope for Alzheimer’s disease therapeutics. Journal of Advanced Research 40, 207–221, doi: 10.1016/j.jare.2021.07.001 (2022).

49 Thapar, N. et al. Application of CRISPR/Cas9 in the management of Alzheimer’s disease and Parkinson’s disease: a review. Ann Med Surg (Lond) 86, 329–335, doi:10.1097/ms9.0000000000001500 (2024).

50 Chacko, L., Chaudhary, A., Singh, B., Dewanjee, S. & Kandimalla, R. CRISPR-Cas9 in Alzheimer’s disease: Therapeutic trends, modalities, and challenges. Drug Discov Today 28, 103652, doi:10.1016/j.drudis.2023.103652 (2023).

51 Kiani, L. New CRISPR-based strategies for Alzheimer disease. Nature Reviews Neurology 19, 507–507, doi:10.1038/s41582-023-00856-5 (2023).

52 Sheridan, C. The world’s first CRISPR therapy is approved: who will receive it? Nat. Biotech. News 42, 3–4, doi: 10.1038/d41587-023-00016-6 (2023).

53 He, S. The first human trial of CRISPR-based cell therapy clears safety concerns as new treatment for late-stage lung cancer. Signal Transduction and Targeted Therapy 5, 168, doi:10.1038/s41392-020-00283-8 (2020).

54 Hampton, T. With First CRISPR Trials, Gene Editing Moves Toward the Clinic. Jama 323, 1537–1539, doi:10.1001/jama.2020.3438 (2020).

55 Gillmore, J. D. et al. CRISPR-Cas9 In Vivo Gene Editing for Transthyretin Amyloidosis. N Engl J Med 385, 493–502, doi:10.1056/NEJMoa2107454 (2021).

56 Henderson, H. CRISPR Clinical Trials: A 2023 Update. Innovative Genomics https://innovativegenomics.org/news/crispr-clinical-trials-2023/ (2023).

57 Taha, E. A., Lee, J. & Hotta, A. Delivery of CRISPR-Cas tools for in vivo genome editing therapy: Trends and challenges. J Control Release 342, 345–361, doi:10.1016/j.jconrel.2022.01.013 (2022).

58 Subbiah, N. et al. Deformable Nanovesicles Synthesized through an Adaptable Microfluidic Platform for Enhanced Localized Transdermal Drug Delivery. J Drug Deliv 2017, 4759839, doi:10.1155/2017/4759839 (2017).

59 Okawa, H. et al. Mechanism of bisphosphonate-related osteonecrosis of the jaw (BRONJ) revealed by targeted removal of legacy bisphosphonate from jawbone using competing inert hydroxymethylene diphosphonate. Elife 11, e76207, doi:10.7554/eLife.76207 (2022).

60 Seo, H. & Lee, H. Recent developments in microfluidic synthesis of artificial cell-like polymersomes and liposomes for functional bioreactors. Biomicrofluidics 15, 021301, doi:10.1063/5.0048441 (2021).

61 Bae, K. H. et al. Durable cross-protective neutralizing antibody responses elicited by lipid nanoparticle-formulated SARS-CoV-2 mRNA vaccines. npj Vaccines 9, 43, doi:10.1038/s41541-024-00835-x (2024).

62 Gaudelli, N. M. et al. Directed evolution of adenine base editors with increased activity and therapeutic application. Nat Biotechnol 38, 892–900, doi:10.1038/s41587-020-0491-6 (2020).

63 Harris, F. M. et al. Astroglial regulation of apolipoprotein E expression in neuronal cells. Implications for Alzheimer’s disease. J Biol Chem 279, 3862–3868, doi:10.1074/jbc.M309475200 (2004).

64 Zhang, M., Guller, S. & Huang, Y. Method to enhance transfection efficiency of cell lines and placental fibroblasts. Placenta 28, 779–782, doi:10.1016/j.placenta.2007.01.012 (2007).

65 Kluesner, M. G. et al. EditR: A Method to Quantify Base Editing from Sanger Sequencing. Crispr j 1, 239–250, doi:10.1089/crispr.2018.0014 (2018).

66 Dupont-Wallois, L. et al. ApoE synthesis in human neuroblastoma cells. Neurobiol Dis 4, 356–364, doi:10.1006/nbdi.1997.0155 (1997).

67 Holly, O. et al. Intraneuronal β-Amyloid Aggregates, Neurodegeneration, and Neuron Loss in Transgenic Mice with Five Familial Alzheimer’s Disease Mutations: Potential Factors in Amyloid Plaque Formation. The Journal of Neuroscience 26, 10129, doi:10.1523/JNEUROSCI.1202-06.2006 (2006).

68 Finn, J. D. et al. A Single Administration of CRISPR/Cas9 Lipid Nanoparticles Achieves Robust and Persistent In Vivo Genome Editing. Cell Rep 22, 2227–2235, doi:10.1016/j.celrep.2018.02.014 (2018).

69 Qiu, M. et al. Lipid nanoparticle-mediated codelivery of Cas9 mRNA and single-guide RNA achieves liver-specific in vivo genome editing of Angptl3. Proceedings of the National Academy of Sciences of the United States of America 118, doi:10.1073/pnas.2020401118 (2021).

70 Yokel, R. A. Nanoparticle brain delivery: a guide to verification methods. Nanomedicine (Lond) 15, 409–432, doi:10.2217/nnm-2019-0169 (2020).

71 Yokel, R. A. Methods to Quantify Nanomaterial Association with, and Distribution Across, the Blood-Brain Barrier In Vivo. Methods in molecular biology (Clifton, N.J.) 1894, 281–299, doi:10.1007/978-1-4939-8916-4_16 (2019).

72 Kim, Y. B. et al. Increasing the genome-targeting scope and precision of base editing with engineered Cas9-cytidine deaminase fusions. Nature Biotechnology 35, 371–376, doi:10.1038/nbt.3803 (2017).

73 Kazemian, P. et al. Lipid-Nanoparticle-Based Delivery of CRISPR/Cas9 Genome-Editing Components. Mol Pharm 19, 1669–1686, doi:10.1021/acs.molpharmaceut.1c00916 (2022).

74 Yao, Z. et al. A high-resolution transcriptomic and spatial atlas of cell types in the whole mouse brain. Nature 624, 317–332 (2023). 10.1038/s41586-023-06812-z

75 Sanabria-Diaz, G., Melie-Garcia, L., Draganski, B., Demonet, J.-F. & Kherif, F. Apolipoprotein E4 effects on topological brain network organization in mild cognitive impairment. Scientific Reports 11, 845, doi:10.1038/s41598-020-80909-7 (2021).

76 Cho, H. et al. Distribution and clinical impact of apolipoprotein E4 in subjective memory impairment and early mild cognitive impairment. Scientific Reports 10, 13365, doi:10.1038/s41598-020-69603-w (2020).

