## Supplementary material for "Successful Gene Editing of Apolipoprotein E4 to E3 in Brain of Alzheimer Model Mice After a Single IV Dose of Synthetic Exosome-Delivered CRISPR": Supplemetary Material

Bruce Teter[1*], Jesus Campagna[1*], Chunni Zhu[1], Grace E. McCauley[2], Patricia Spilman[1], Donald B. Kohn[2], Varghese John[1§]
[1]The Drug Discovery Lab (DDL), Mary S. Easton Center for Alzheimer's Disease Research and Care, Department of Neurology, Geffen School of Medicine, UCLA, Los Angeles, CA
[2] Kohn Lab, Department of Microbiology, Immunology and Genetics, UCLA, Los Angeles, CA

**Supplementary Figures**

**Fig. S1.** Particle size analysis of SE-CRISPR
**Fig. S2.** Integrity analysis of CRISPR RNAs
**Fig. S3**. Brain Capillary Depletion Acid Phosphatase assay

**Supplementary Tables**

**Table S1.** Tested combinations of gRNA and CBR mRNA
**Table S2.** E4 to E3 Editing E3%
**Table S3.** E4 to E3 Editing E3% in brain fractions

### Supplementary Figures

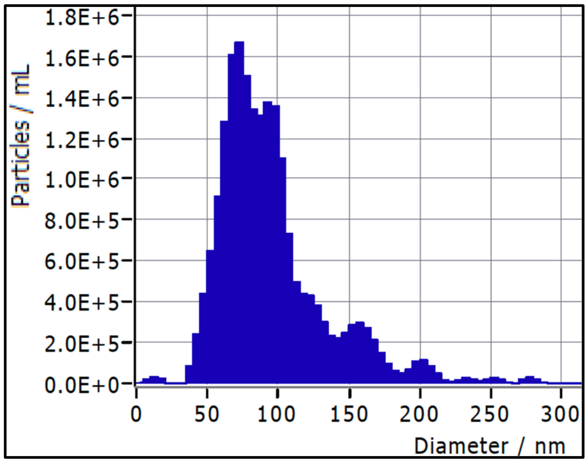


**Fig. S1.** *Particle size analysis of SE-CRISPR*. Nanotracker analysis of Exoglo-labeled vesicles indicate an average diameter of 90 nm**.**

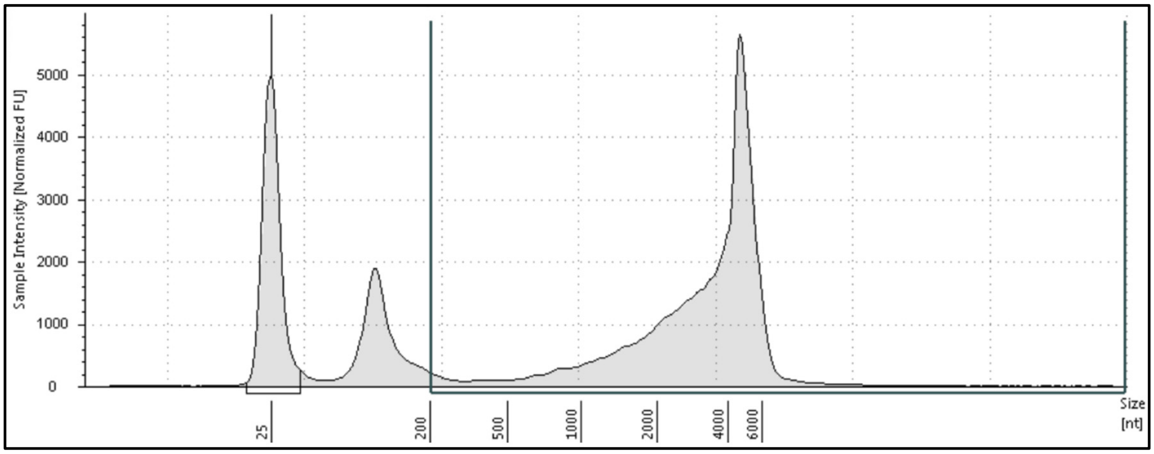


**Fig. S2**. *Integrity analysis of CRISPR RNAs*. Tapestation analysis of CRIPSR gRNA (100nt) and CBE mRNA (5940nt) reveals that after encapsulation into SEs and subsequent retrieval from SEs, gRNA and CBE mRNA are intact and of the expected sizes. The peak at 25 nt is an exogenous RNA marker.

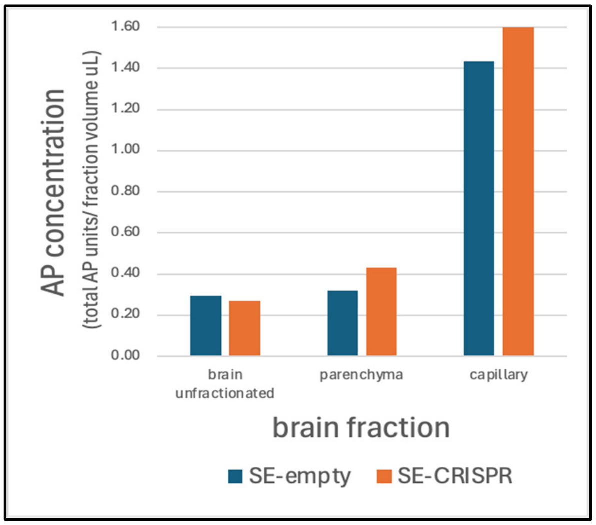


**Fig. S3.** *Brain Capillary Depletion Acid Phosphatase assay*. Mouse brain hemispheres were fractionated by homogenization and centrifugation in dextran. Acid phosphatase (AP) was measured in unfractionated and the dextran centrifugation fractions parenchyma (dextran supernatant) and capillary (dextran pellet). The AP concentration in each fraction as calculated by total AP units in the fraction, divided by the fraction's volume in µL (total AP units/ fraction volume µL). Results are shown for the brains used for E3 editing measurement, one from a mouse treated with CRISPR-containing SE (SE-CRISPR) and the other treated with empty SE (SE-empty).

### Supplementary Tables

| **Table S1**. Tested combinations of gRNA and CBE mRNA | | | |
|---|---|---|---|
| | Combination | | E3% editing* |
| 1 | gRNA #1 | pBT375 | 14 |
| 2 | gRNA #2 | pBT375 | 51-54 |
| 3 | gRNA #1 | pYE1BE4max | <7 |
| 4 | gRNA #2 | pYE1BE4max | <7 |
| gRNA #1 - GGACGTGCGCGGCCGCCTGGTG<br>gRNA #2 - CGTGCGCGGCCGCCTGGTGCTG | | | *Measure by EditR |

| **Table S2.** E4 to E3 Editing E3% | | | |
|---|---|---|---|
| | **Liver** | **Brain** | **Buffy Coat** |
| **SE-Empty** | 0.0057 | 0.0058 | 0.0162 |
| **SE-CRISPR** | 0.809 | 0.141 | 0.357 |
| **Normalized to SE-Empty** | 0.803 | 0.135 | 0.341 |

| **Table S3.** E4 to E3 Editing E3% in brain fractions | | | |
|---|---|---|---|
| | unfractionated | parenchyma | capillary |
| **%E3 editing** | 0.049 | 0.035 | 0.007 |
